# Extracellular Vesicles Carry Distinct Proteo-Transcriptomic Signatures That are Different from Their Cancer Cell of Origin

**DOI:** 10.1101/2021.09.20.460963

**Authors:** Tzu-Yi Chen, Edgar Gonzalez-Kozlova, Taliah Soleymani, Sabrina La Salvia, Natasha Kyprianou, Susmita Sahoo, Ashutosh K. Tewari, Carlos Cordon-Cardo, Gustavo Stolovitzky, Navneet Dogra

## Abstract

Circulating extracellular vesicles (EVs) contain molecular footprints from their cell of origin and may provide potential non-invasive access for detection, characterization, and monitoring of numerous diseases. Despite their growing promise, the integrated proteo-transcriptomic landscape of EVs and their donor cells remain poorly understood. To assess their cargo, we conducted small RNA sequencing and mass spectrometry (LC-MS/MS) of EVs isolated from *in vitro* cancer cell culture and prostate cancer patients’ serum. Here, we report that EVs enrich for distinct molecular cargo, and their proteo-transcriptome is predominantly different from their cancer cell of origin, implicating a coordinated disposal and delivery mechanism. We have discovered that EVs package their cargo in a non-random fusion, as their most enriched RNAs and proteins are not the most abundant cargo from their donor cells. We show that EVs enrich for 4 times more cytoskeletal and 2 times extracellular proteins than their donor cells. While the donor cells carry 10 times more mitochondrial and 3 times nuclear proteins than their EVs. EVs predominantly (40-60%) enrich for small RNA (~15-200 nucleotides) molecules that implicate cell differentiation, development, and signaling signatures. Finally, our integrated proteo-transcriptomic analyses reveal that EVs are enriched of RNAs (RNY3, vtRNA, and MIRLET-7) and their complementary proteins (YBX1, IGF2BP2, SRSF1/2), implicating an interrelated mechanism that may protect and regulate transcripts until a biological function is achieved. Based on these results, we envision that the next-generation clinical assays will take an integrative multi-omic (proteomic and transcriptomic) approach for liquid biopsy in numerous diseases.

## Introduction

Emerging evidence suggests that circulating EVs contain molecular footprints - lipids, proteins, metabolites, RNA, and DNA - from their cell of origin for intercellular communication as well as cargo delivery and disposal(Kowal et al., 2014; Simpson et al., 2009; Valadi et al., 2007). Consequently, EVs are increasingly recognized as key players in the signal transduction, priming of tumor microenvironment, and metastasis(Archer et al., 2020; N. Dogra et al., 2020), mainly through cellular crosstalk and vesicle trafficking(Archer et al., 2020; Hoshino et al., 2015; Kamerkar et al., 2017). First described in the 1980s(Johnstone et al., 1987; Trams et al., 1981), exosomes are extracellular nanovesicles of endocytic origin, which are formed by the inward budding of a late endosome, also known as multivesicular body (MVB)(Kowal et al., 2014; Raposo and Stoorvogel, 2013; Simpson et al., 2009). Subsequently, MVBs fuse with the plasma membrane resulting in the release of exosomes to the extracellular environment(Johnstone et al., 1987; Trams et al., 1981). Nevertheless, exosomes are merely a subset of secretory EVs, which include – but are not limited to - apoptotic, micro-, and onco-vesicles(Kalluri and LeBleu, 2020; Kowal et al., 2014; Raposo and Stoorvogel, 2013). Accumulating evidence shows that while all EVs carry molecular footprints from their cell of origin; exosomes selectively package proteins, nucleic acids, and do not appear enriched in cellular debris(Kowal et al., 2014; Lotvall et al., 2014). In understanding the complete cargo of EVs, recent studies have successfully utilized proteomic and transcriptomic technologies for their molecular analyses and multiple databases have compiled their outcomes accordingly(Bellingham et al., 2012; Murillo et al., 2019; Simpson et al., 2009). “Extracellular RNA (exRNA) Atlas” and “Vesiclepedia” are two of such comprehensive databases(Kalra et al., 2012; Murillo et al., 2019; Rozowsky et al., 2019).

Currently, majority of the published EV literature are analyses of either the proteomic or the transcriptomic signatures alone(Keerthikumar et al., 2015; Mathivanan et al., 2010; Simpson et al., 2009). While, numerous studies have investigated donor cells and their EVs’ content, whether their proteins and RNAs are functionally interrelated, remain poorly understood. This is mainly due to a lack of proteomic and transcriptomic analyses of matching donor cells and their EVs. An integrative proteo-transcriptomic analysis of EVs will not only reveal the mutual regulation between RNA and proteins, but will also provide key insights into their applications as therapeutic and non-invasive next-gen liquid-biopsy procedures(Archer et al., 2020; Smith et al., 2018).

In this study, we have performed a comprehensive proteomic and transcriptomic analysis of EVs derived from *in vitro* cancer cell culture and prostate cancer patient’s serum (Fig. 1). To identify EV specific molecular signature, we compared the RNA and protein profile of EVs with their donor cells and assigned to their respective subcellular locations according to Gene Ontology (GO) annotation. Next, we asked if the EV’s protein and RNA cargo are inter-related and converge to achieve the same biological function. To address this, we utilized RNA-Interactome of the protein and RNA cargo of EVs and examined for functionality, mutual regulations, and distinct cellular pathways. Finally, we put forth a comprehensive model that integrates proteomic and transcriptomic signature of EVs, which will provide a conceptual advance in development of next-generation clinical assays by taking a multiomic (proteomic and transcriptomic) approach for liquid biopsy in numerous diseases.

**Figure 1.**
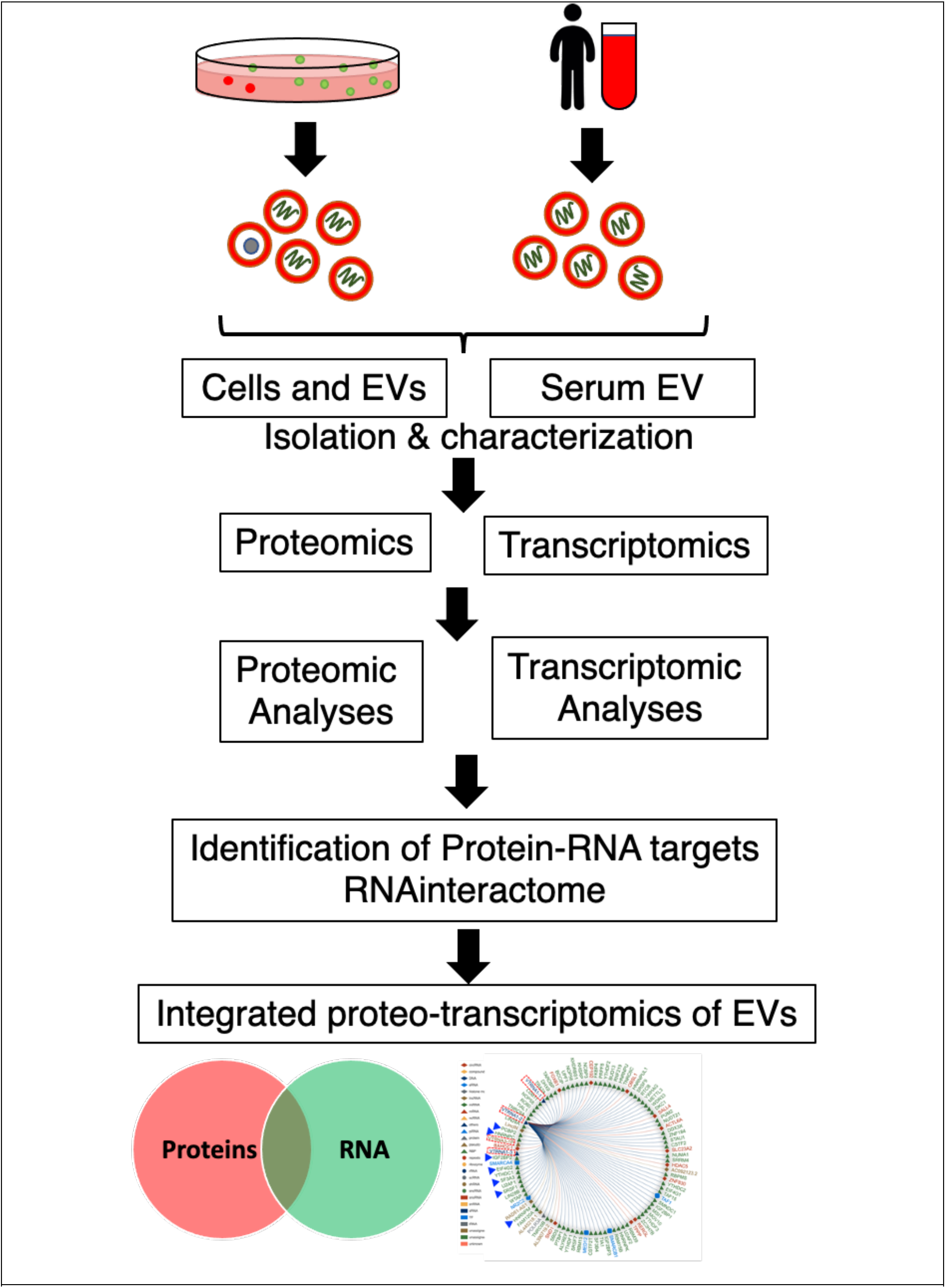
Overview of experimental design and data analysis. Extracellular vesicles (EVs) are isolated from prostate cancer cell culture media and patient serum. Subsequently, the proteo-transcriptome of EVs is characterized.

## Results

Our study has generated three datasets of proteomics and transcriptomics from the donor cancer cells, their EVs, and prostate cancer patient’s serum EVs (Fig. 1). In total, we have studied ~20 proteomic and ~22 transcriptomic profiles. The three datasets were labeled as: **1**) A “*transcriptomics*” dataset (n=6) composed of small RNAseq of donor cancer cells and their EVs. **2**) A “*proteomics*” dataset (n=15) composed of mass spectrometric analyses of donor cancer cells and their EVs. **3**) A distinct “*human serum EV*” cohort (n=6) composed of proteomics and transcriptomics for prostate cancer patient serum EVs.

### Characterization of extracellular vesicles

To characterize the size, shape, morphology, and canonical markers of EVs, we conducted their comprehensive quality control analysis. These analyses included transmission electron microscopy (TEM), immuno-gold TEM, Nanoparticle Tracking Analyses (NTA), zeta potential analyses, and multi-color immuno-fluorescence colocalization analyses (nanoview). First, to characterize their morphology, donor cells and their EVs were examined under a high-resolution TEM revealing the release of EVs from the donor cells that illustrated a typical cup-shaped morphology of the EVs (Fig. 2A). We observed two mechanisms of EV secretion; (1) vesicles of endocytic origin (exosomes) secreted via fusion of MVBs with the plasma membrane; and (2) secretion via budding (exocytosis) from the plasma membrane. Notably, the first mechanism displayed consistent round and cup-shaped ~80 nm vesicles, while the second mechanism displayed relatively larger and heterogenous, ~50-250 nm vesicles. Then, the EVs were isolated from the cell culture media. The isolated EVs were observed under a TEM revealing vesicles of ~80-100 nm sizes (Fig. 2B). The Immuno-gold TEM captured CD81 (a canonical exosome marker) antibody attached to 6 nm colloidal gold particles (Fig. 2C). Further, immuno-fluorescence colocalization analyses (see methods) of EVs revealed three vastly studied canonical tetraspanin exosome markers CD81, CD9, and CD63 on vesicles surface (Fig. 2D). As expected, all three exosomal tetraspanins were enriched on the vesicle’s surface, confirming the presence of canonical markers for exosomes. In contrast, the negative control mouse IgG was non-specific to the EVs and displayed no signal for tetraspanin proteins (Fig. 2D & E bottom). The EVs’ zeta potential ranged from −30 to +30 mV (highest at −3 mV) demonstrating moderate colloidal stability between the EVs and their surrounding fluid environment (Fig. 2G). Taken together, the isolated EVs carry canonical tetraspanins for exosomes, are between the size of ~50 to 200 nm in diameter and the majority of the EVs enriched for a median of ~80 nm sized particles, demonstrating moderate colloidal stability (Fig. 2B, C, F).

**Figure 2.**
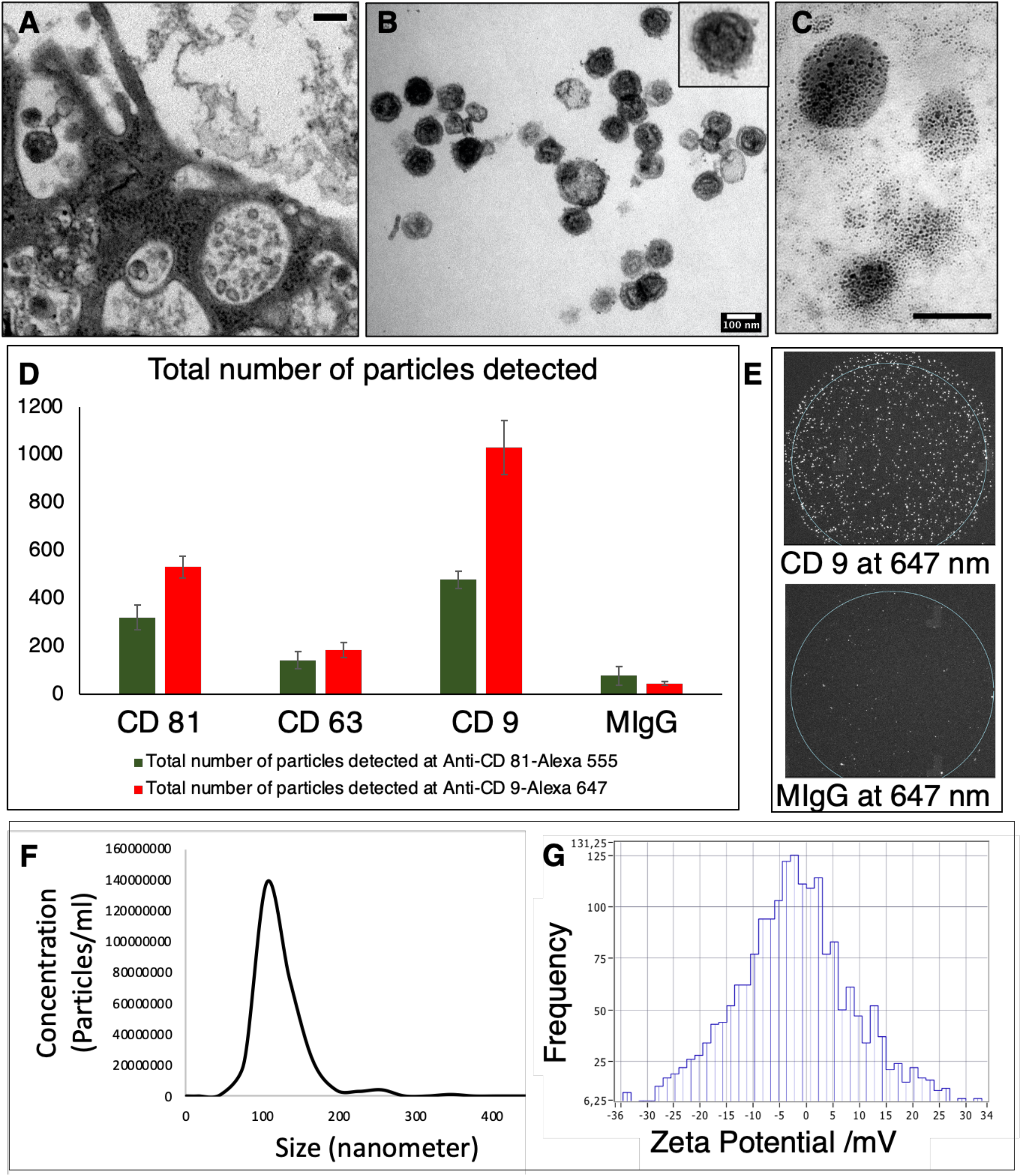
Characterization Serum and cell culture derived EVs. (A) TEM image of EVs released from cells and isolated EVs (B). (C) canonical markers Immunogold (6nm) labeled CD81 shows specificity to the vesicle surface. Scale bar is 100nm. (D) immuno-fluorescence co-localization analyses of EVs confirmed presence of canonical tetraspanins on vesicle surface. Mouse IgG antibody is a negative control. (E) Optical image of immuno-fluorescence co-localization analyses shows EVs specificity to CD9 but not to mouse IgG. (F) A size and concentration analyses of isolated EVs show distribution of vesicles (50-250 nm). (G) Zeta potential of EVs show moderate colloidal stability between the EVs and their surrounding fluid environment.

### Characterization of the proteome of EVs and donor cells

To characterize the proteomic landscape of donor cells and their EVs, we extracted total protein, conducted mass spectrometry (LC-MS/MS), and investigated their respective protein datasets. To reliably identify a protein, positive identification was set at 5% protein false discovery rate (FDR) and 1% peptide FDR. All samples were analyzed in duplicates. As a result, 422 proteins for EVs and 5630 proteins for cells passed our filtering criteria. Notably, the top 5% of EV proteins were molecules of endosomal sorting complex required for transport (ESCRT), endosomal origin (clathrin), multivesicular body (MVB12A), membrane trafficking (RAB proteins, annexins), cytoskeletal (actin, tubulin, myosin), heat shock proteins (HSC70, HSP71, HSP90), and exosomal markers of endocytic origin (ALIX), and TSG101. These findings are in accordance with literature, as ESCRT and other endocytic machinery proteins are enriched in exosomes(Kalra et al., 2012; Simpson et al., 2009).

Next, we utilized the large proteomic dataset from Vesiclepedia database, which is a compilation of data from over ~1300 studies from EVs(Kalra et al., 2012). A comparison of our proteomic data of EVs with Vesiclepedia database displayed that ~94% (396 proteins) of the EVs proteins in our study were present in the Vesiclepedia, while only ~6% (26) proteins were unique to our EVs study (Fig. 3A). These results confirmed that our EV isolation methods and proteomic analyses were consistent, reproducible, and reliable with respect to the other ~1300 studies present in the Vesiclepedia database.

**Figure 3.**
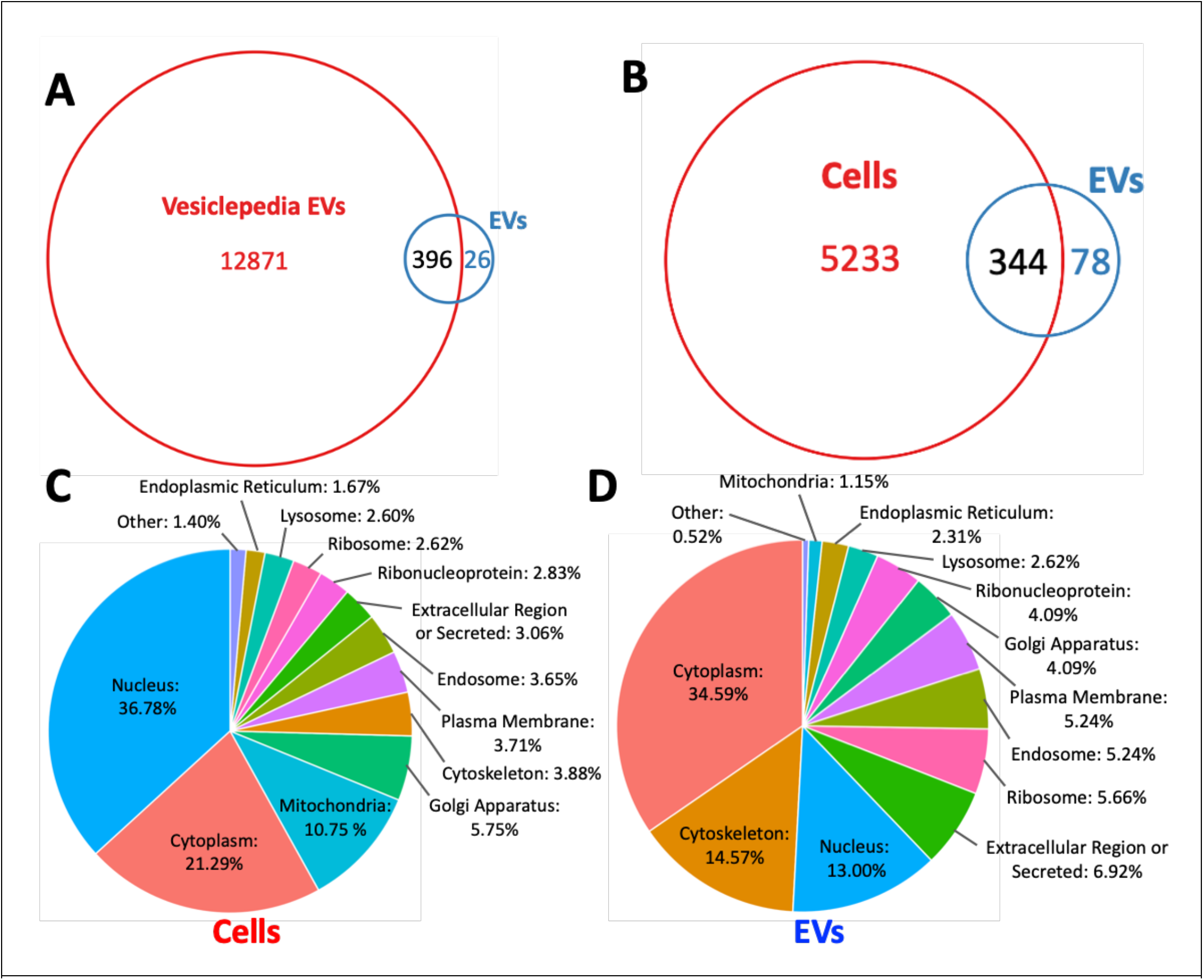
Proteomic analysis of EVs and their donor cells. (A) Comparative analyses of Vesiclepedia proteomic database of EVs with in-house proteomics of EVs isolated from prostate cancer cell lines, 22RV1. (B) Comparative analyses of EV and their donor cell proteins. (C) Subcellular location of donor cell proteome. (D) Subcellular location of EVs proteome.

Finally, we compared the proteomics of donor cells with their EVs (Fig. 3B). Of note, ~22% (78) of the proteins were unique to the EVs, while ~78% (344) of the EVs proteins displayed an overlap with their donor cell proteins. To compare the subcellular location, both donor cell and EV proteins were assigned to their respective locations according to Gene Ontology (GO) annotation (Fig. 3C & D). These analyses revealed that cells primarily contain nuclear proteins (37% vs 13% in EVs), while EVs contain cytoplasmic proteins (35% vs 21% in cells). EVs were enriched for 4 times more cytoskeletal and 2 times more extracellular proteins than their donor cells. While the donor cells carry 10 times more mitochondrial and 3 times nuclear proteins than their EVs. Overall, the proteomic profile of EVs is distinct with many proteins specifically enriched in EVs but not in their donor cells, implicating an unexplained mechanism that may stem from a coordinated effort for cell-to-cell communication, disposal, and delivery processes(Kalra et al., 2012; Simpson et al., 2009).

### Characterization of the transcriptome of EVs and donor cells

Recent studies have shown that EVs contain diverse small RNA subtypes and their sizes range between ~20-200 nt(Srinivasan et al., 2019; Wei et al., 2017). However, the type of RNA packaged inside EVs remain a matter of intense debate(Srinivasan et al., 2019; Valadi et al., 2007; Wei et al., 2017). To assess and compare the precise length of RNA of donor cells and their EVs, we extracted total RNA and characterized through capillary electrophoresis using two separate analyses kits: Pico and small RNA kits for bioanalyzer (Fig. 4A & Sup. Fig. 1). These analyses revealed that although both donor cells and their EVs displayed small RNAs, their lengths are different. For instance, a major peak around ~100 nt is predominantly present in EVs, while cells contain additional small RNA peaks at ~80 nt and ~50 nt (Fig. 4A). EVs contain major peaks at ~60 and 100 nt, which may correspond to tRNA and small nuclear RNA (snRNA) (Sup. fig. 1). These observations compelled us to survey the current small RNA landscape that range below 200nt(Murillo et al., 2019; Wei et al., 2017). Among the small RNAs, miRNAs (~21 nt), siRNAs (~20-25bp), tRNAs (~60-95) nt, 5S rRNAs (~120 nt), and snRNAs (~150 nt) are potential candidates that maybe present in EVs(Boivin et al., 2019). To address the different subtypes of RNAs in EVs and cells, we proceeded with cDNA library preparation followed by total small RNA sequencing. A detailed analysis of the transcriptomic profiles is discussed in detail below.

**Fig 4.**
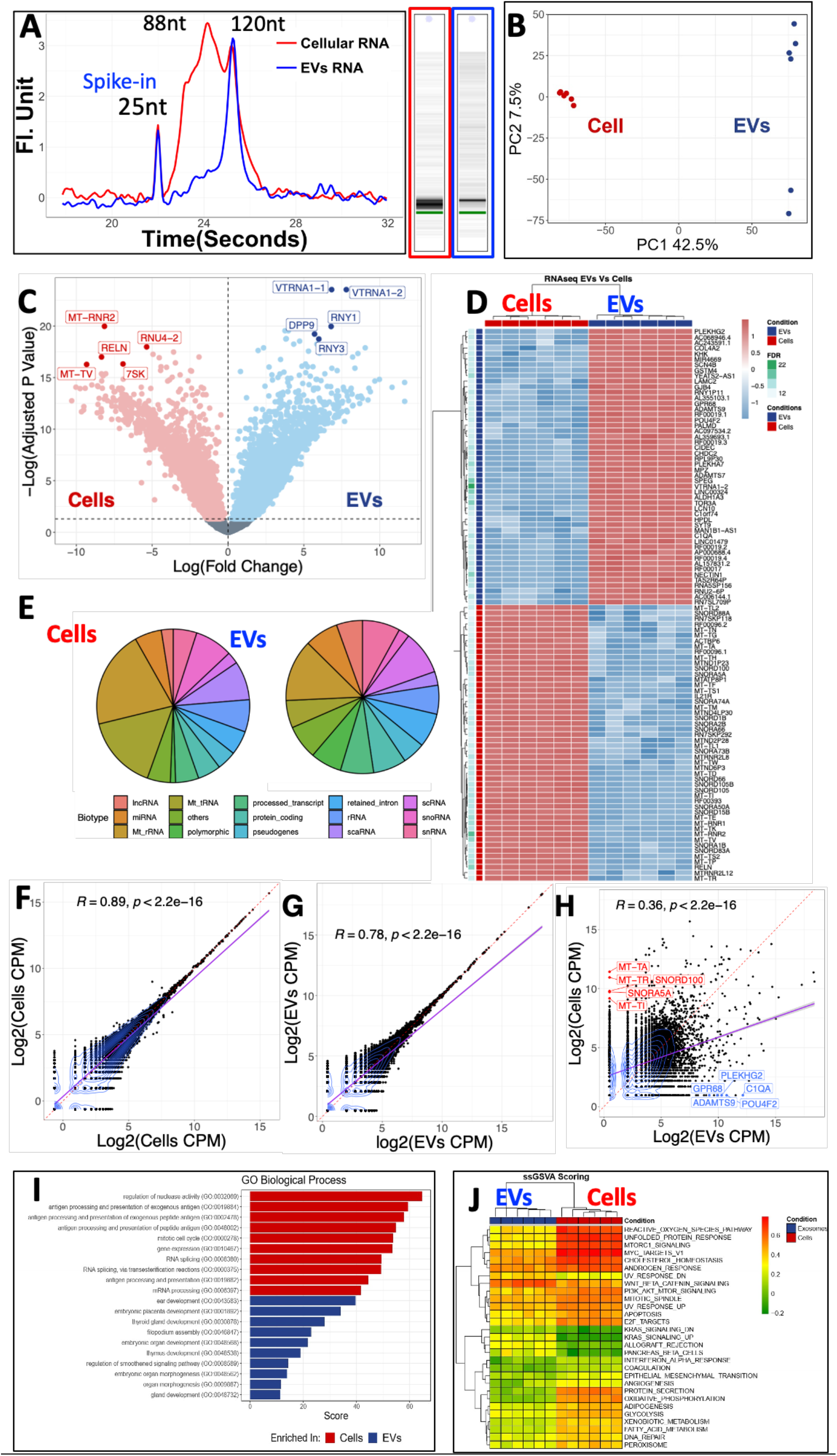
Transcriptomic analyses of EVs and donor cells. (A) Capillary electrophoresis analyses of small RNA from donor cells and their EVs. (B) PCA of the RNA content between cells and EVs. (C) Volcano plot between Cells and EVs. (D) Differentially expressed RNAs between Cells and EVs. (E) Differences between gene biotypes detected between cells and EVs. Reproducibility study of cell versus cell RNAseq (F), EVs versus EVs (G) and comparison cells versus their EVs RNA (H). (I) Top enriched pathways for EVs and Cells. G. Hallmark cancer pathways identified in EV and cellular RNA.

### EVs enrich for unique RNA signatures that are different from their cell of origin

Although there are no established canonical RNA markers for EVs, a recent extracellular RNA (exRNA) study investigated and compared exRNA cargo types in over 5000 human samples using different isolation methods and biofluids. This independent analysis found that our isolation methods (UC and nanoDLD chip technology(Smith et al., 2018)) specifically isolate low- and high-density vesicles (cargo type 1 & 4, respectively) with minimum contamination from the lipoproteins and argonaute proteins(Murillo et al., 2019). Here, to identify the entire payload of the donor cells and their EVs’ RNA, we conducted total small RNA sequencing and investigated their gene expression profiles (Fig. 4B, C, D, & E)). We identified over ~20, 000 distinct RNA molecules in EVs and donor cells. The composition of RNA molecules in EVs was composed of 45% proteincoding, 65% non-coding with a dominant subpopulation of 25% miRNAs, followed by tRNA, snRNAs, snoRNAs, and lncRNAs each under 4% (Fig. 4E). Next, we performed principal component analysis (PCA) showing that the molecular profiles were uniquely different between cells and EVs with a total of 50% variance explained by PC1 and PC2 (Fig. 4B). The variance between EVs and their cell of origin indicates that the most abundant transcripts in cells were different from those in EVs. To understand the landscape of RNA within cells and their EVs, we performed a differential expression analysis using mixed linear models revealing over 8000 differentially expressed RNAs with a log fold change up to 10 times (Fig. 4C & D). The observations of gene biotype revealed that distinct cargo types are loaded into EVs. While EVs are packaged with effector and regulatory RNA molecules, cells predominantly contain mtRNAs and rRNAs (Fig. 4D).

Our RNAseq studies were reproducible as replicates of cell versus cell (from two separate *in vitro* cell cultures) and EVs versus EVs yielded Rho = ~0.9 and ~0.8, respectively (Fig. 4F & G). In contrast, the correlation between donor cells versus their EVs yielded a low correlation with Rho = 0.3 (Fig. 4H). Interestingly, numerous RNA molecules are exclusively detected in the EVs (Fig. 4H). EVs exclusively contain ADAMTS9, PLEKHG2 (nucleotide binding protein), GPR68 (a proton sensing G protein-coupled receptor 68), POU4F2, and Let-7 microRNA precursor, which are involved in cell development, signal transduction, and cancer metastasis(Schaefer et al., 2009; Shurtleff et al., 2017; Valadi et al., 2007). On the other hand, donor cells exclusively contained mitochondrial RNA and small nucleolar RNA (SNORA 5A and SNORD100). Consistent with these observations, pathway enrichment analysis revealed that EVs are enriched in RNA molecules that implicate cell differentiation, development, and morphogenesis signatures (Fig. 4I). In contrast, cells enrich from pathways associated with cellular and metabolic processes. Additionally, the results of gene set enrichment analyses of hallmark cancer pathways showed that cells were enriched for cellular damage response and that these signatures were retained in EVs for pathways such as MYC targets or androgen response(Gross et al., 2012; Miyamoto et al., 2015; Zhang and Wrana, 2014). Wnt beta-catenin and KRAS signaling was enhanced in EVs, suggesting EVs involvement in promoting cancer progression and metastasis (Fig. 4J). This observation is in accordance with multiple studies demonstrating the significance of Wnt pathways in cell regulatory processes including cell proliferation, stem cell differentiation, and migration(Murillo et al., 2019; Smith et al., 2018). Taken together, these results suggest that the RNA of donor cells and EVs are distinct and EVs primarily enrich for specific small RNA that target cell differentiation, development, and signaling processes.

### Proteo-transcriptomic analyses of EVs from prostate cancer patient serum reveal their mRNA binding and immune regulation function

The goal of this pilot proteo-transcriptomic analysis was to complement our *in vitro* studies with human serum-derived EVs. Encouraged by our studies from cells and EVs, we isolated EVs from the serum of 6 prostate cancer patients and investigated their proteo-transcriptome. To understand EVs’ content irrespective of the isolation method used(Murillo et al., 2019), we used two separate technologies (buoyant density based UC, and size based nanoDLD(N. Dogra et al., 2020; Smith et al., 2018)) to isolate EVs followed by their proteo-transcriptomic analyses. First, we characterized serum EVs for their size, shape, morphology, and canonical marker of EVs. Similar to cell culture derived EVs, TEM showed round, cup shaped morphology for serum EVs (Fig 5A). The EVs ranged from ~50 to 150nm in size and carried CD9 on their surface (Fig. 5A, B, C). Rigorous characterization of UC and nanoDLD isolated EVs from various biofluids has been presented in our previous publications(Murillo et al., 2019; Smith et al., 2018). Our proteo-transcriptomic analyses from serum EVs displayed specific signals and highlighting the presence of an EV specific signature (Fig. 5D, E). The serum EV signatures from RNAseq analysis are associated with protein binding, epithelial cell proliferation, post-translational protein regulation, and regulation of synaptic plasticity (Fig. 6A). The pathway enrichment results of the mass spectrometry are highly correlated with the regulation of complement cascade or immune response, and processes (Fig. 6B). Overall, similar to the in vitro cell culture, serum EVs carry molecular signatures that implicate protein and RNA regulation.

**Fig 5.**
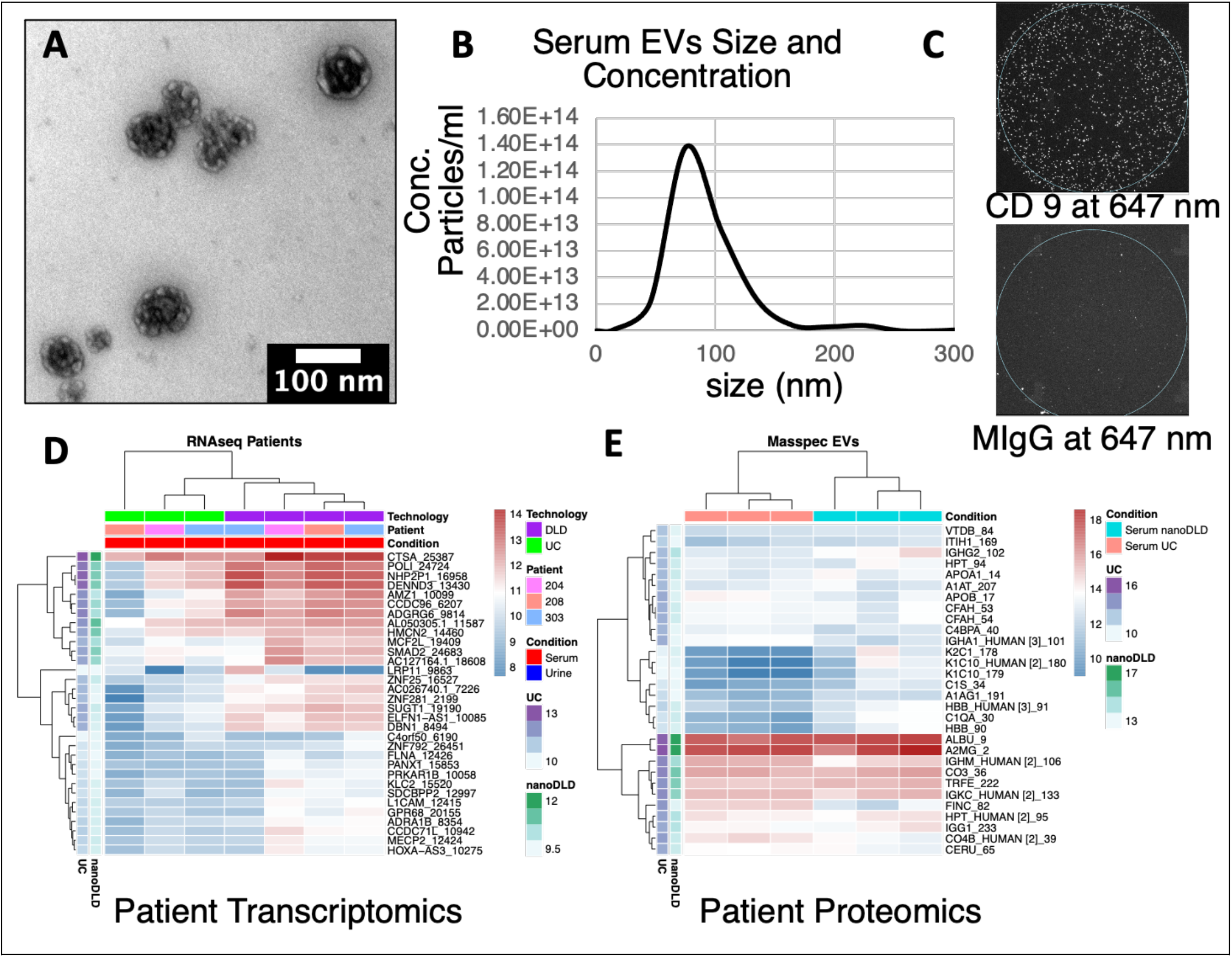
Proteomic & transcriptomic analyses of patient serum EVs. (A) TEM image of UC isolated EVs (B) A size and concentration analyses of serum EVs show distribution of vesicles (50-150 nm). (C) immunofluorescence co-localization analyses serum of EVs confirmed presence of canonical tetraspanin CD9 on vesicle surface. Mouse IgG antibody is a negative control. (D) Top 30 proteins detected in serum EVs. (E) Top 30 RNA detected in serum EVs.

**Fig 6.**
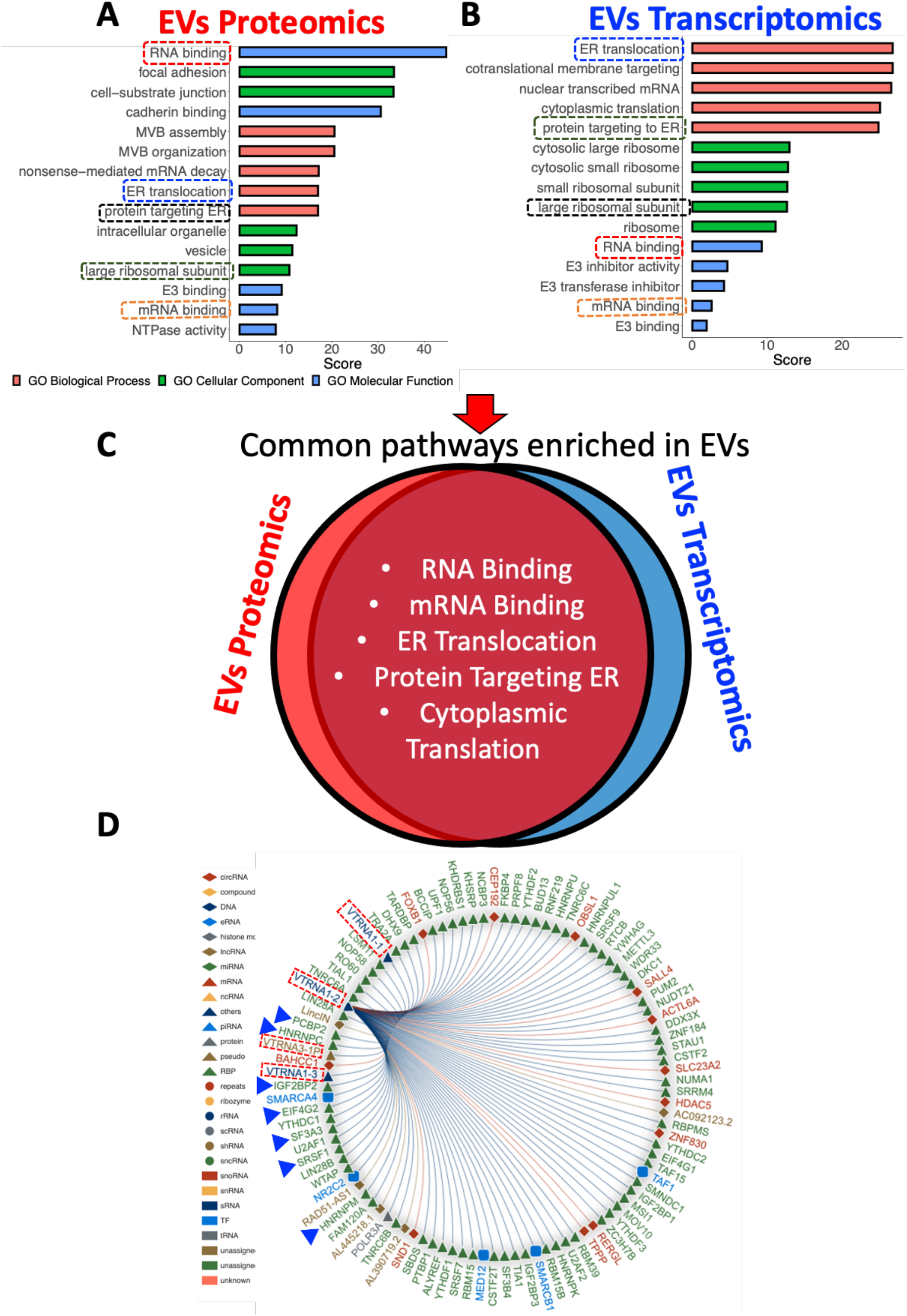
Integrative Proteo-transcriptomic analyses of EVs. Top enriched pathways identified in proteomic (A) and transcriptomic analyses (B) in EV. (C) Overlapping proteo-transcriptomic pathways enriched in EVs. (D) RNAInteractome analyses shows RNA (red box) and their target proteins identified in EVs.

### Integrated proteo-transcriptomic analyses of EVs reveal enrichment of RNA-proteins and RNA-RNA complexes

The amino acids and their complementary nucleotides form protein-RNA complexes that protect and regulate transcripts through their life cycle until a function is achieved(Corley et al., 2020). However, the current landscape of such RNA-protein complexes and their inter-relationship in EVs is poorly understood. Given our comprehensive proteomic and transcriptomic data, we asked whether EVs carry proteins and their complementary RNA. We used RNA interactome database (http://www.rna-society.org/rnainter) to identify EV enriched RNA/proteins and their target RNA-protein and RNA-RNA interactions(Lin et al., 2020).

Here, we discovered that transcription factor YBX1 and its target RNA (RNY3, a Y RNA class) were enriched in our proteo-transcriptomic analyses of EVs (sup. table 1). Vault RNAs (vtRNA 1-1/2/3) were enriched in EVs, and their complementary binding proteins (IGF2BP2, SRSF1) were enriched in our proteomic analyses (Fig. 6D). We also identified various RNA-RNA complexes in EVs. We found that Let-7 (a predominant RNA in EVs) and its associated RNA (SPEG) and protein (SRSF2) were enriched in EVs. Finally, functional annotation analyses revealed that RNA binding, protein translation, and gene expression are the major overlapping molecular pathways between the EV’s RNA and proteins, implicating a coordinated mechanism between EVs and their donor cells (Fig. 6C).

## Discussion

This study provides a conceptual advance in understanding the proteo-transcriptome of EVs from cancer cell culture and human serum (Fig. 7). First, we show that EVs enrich for distinct RNA and protein cargo compared to their cancer cell of origin. We present that EVs enrich for ESCRT, endosomal, multivesicular body proteins, membrane trafficking, and exosomal marker proteins. The cellular sub compartment analyses reveal that cells cargo 10 times more mitochondrial proteins and 3 times more nuclear proteins than EVs. On the other hand, EV cargo 4 times more cytoskeletal, 2 times more extracellular, 1.6 times more cytoplasmic proteins. The RNA characterization revealed that EVs carry a range of small RNA between ~13-200 nt, which display miRNAs, siRNAs, tRNA, 5S rRNAs, and snRNAs. Additionally, considering that the diameter of EVs (~50-150 nm in our studies) allow packaging of RNA up to a limited size, large rRNAs (greater than ~2000 bp) are likely beyond the limits of EVs’ capacity(Valadi et al., 2007; Wei et al., 2017). This observation is supported by a recent study on glioblastoma derived EVs, which showed that ~3000 nt RNA were not present in the small EVs but were abundant in the cells and microvesicles(Wei et al., 2017).

**Fig 7.**
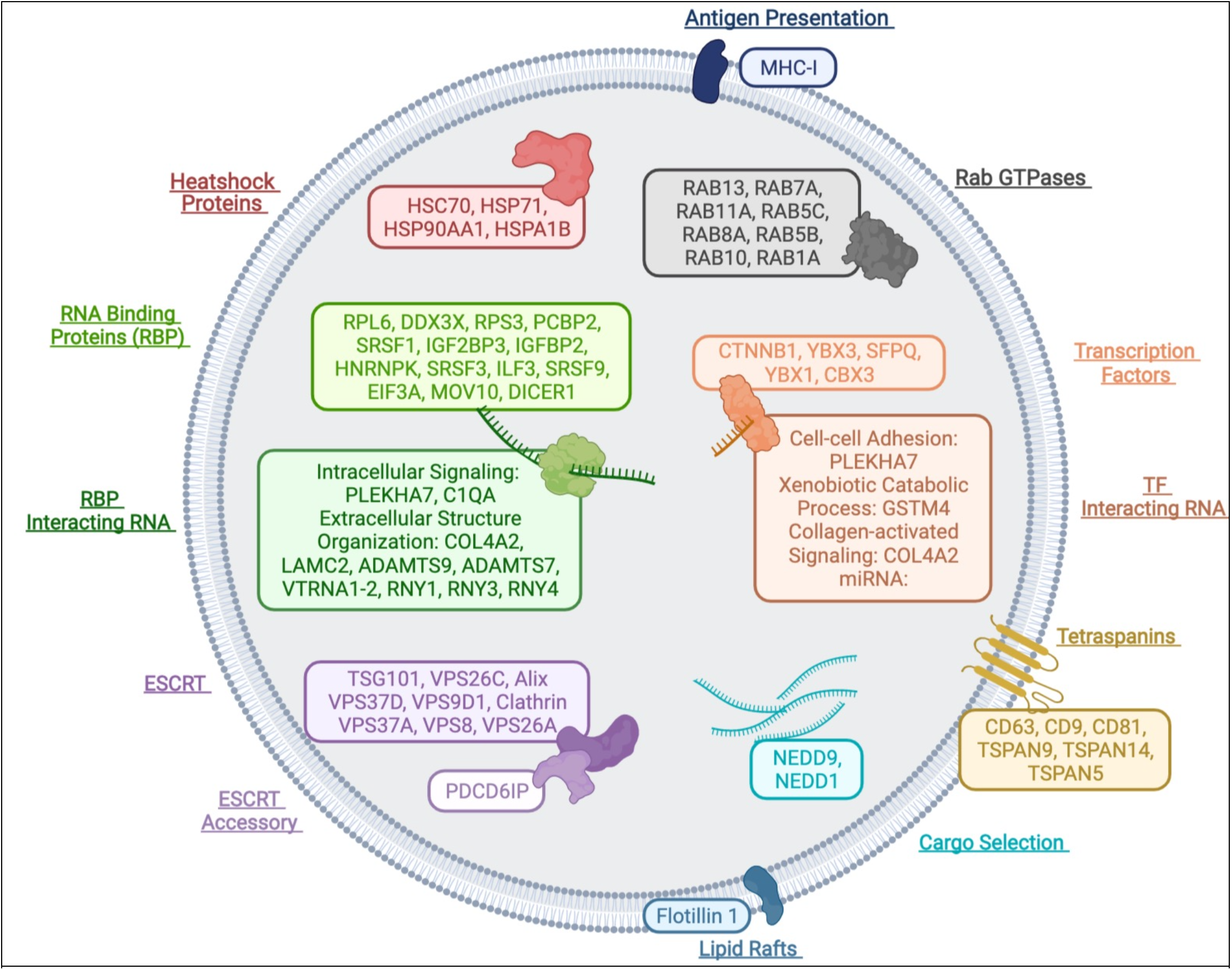
A graphical representation of integrative proteo-transcriptomic composition EVs. Top enriched proteins and their target RNA are shown.

The rational of enrichment of specific proteins and RNA in the EVs (but not in their donor cells) can be explained via following mechanisms: 1) The structural integrity of RNA and proteins inside EVs maybe enhanced due to RNA-RNA or RNA-protein complex formation, while separating them from cytoplasmic nucleases/proteases, hence enhancing their half-life and overall stability. Remarkably, these results also support the hypothesis that oncogenic proteins and RNA in circulating EVs may play a critical role in the dissemination of tumor-associated molecules and could be a viable target for liquid biopsy biomarkers and therapeutic approaches; 2) The delivery and disposal aspect of EVs may implicate a coordinated effort of enrichment of molecules in the EVs while lowering their abundance in the donor cells; 3) The mechanism of packaging of proteins in the secretory vesicles before they leave their donor cells may stem from the cell-to-cell communication, followed by their dissemination mechanism mediated by the EVs.

### Significance of EVs enriched proteins and RNA

The observation that EVs enrich for different molecules then the most abundant cargo in their cell of origin has immense significance with respect to mass EV production for gene delivery and other therapeutic applications. Currently, several industrial and academic institutions are attempting to overexpress RNAs and proteins of interest in donor cells with an assumption that secreted EVs will randomly enrich for the most abundant molecule. However, based on our studies, we find that the most abundant molecules in the cells may not enrich in EVs. These findings suggests that alternate strategies, similar to liposomes encapsulation(Dogra et al., 2019; Dogra et al., 2015; Dogra et al., 2016), must be followed for enriching cargo in the EVs(Dogra et al., 2019; Haraszti et al., 2018).

### Significance of EVs enriched RNA-proteins complexes

We show that EVs enrich for proteins and RNA that are functionally interrelated. These findings are in accordance with previously published studies that have shown intracellular association of several transcription factors, other proteins, and RNA. For instance, RNY3 interacts with transcription factor YBX1, which helps block access to the RNA(Corley et al., 2020; Lin et al., 2020; Shurtleff et al., 2017). A recent study showed YBX1 and its association with mir-223 in exosomes(Shurtleff et al., 2017). We show that enrichment of various RNAs (RNY3, vtRNA, and MIRLET-7) and their complementary proteins (YBX1, IGF2BP2, SRSF1/2) in EVs may target distinct cellular pathways that converge to achieve the same biological goals. Furthermore, these analyses reveal that RNA-protein complex could provide a potential functional interaction network inside EVs to protect and regulate access to the RNA. Thus, EV’s mRNA, miRNA, and proteins may be connected to achieve the same regulatory function.

## Conclusion

In summary, the present study shows that EVs enrich for distinct RNA and protein cargo compared to their cell of origin. This work enables for the first time an integrative proteo-transcriptomic platform from patient serum EVs and their correlation could provide major insight on liquid biopsy and diagnosis of numerous diseases. This is of major clinical significance in the potential use of integrative proteo-transcriptomic platform that would further enhance the diagnostic performance of diseases via liquid biopsy.

## Limitations

We acknowledge the limitation of this study, including low number of patient samples. Despite these limitations, we demonstrate a proof of the principal study and ability to conduct a reproducible and integrative proteomic-transcriptomic analyses on cancer cells’ and patients’ blood derived EVs.

## Author contribution

Conceptualization: ND, GS. Writing - original draft preparation: ND, TC, EGK, TS. Writing - review & editing: TC, EGK, TS, SL, NK, SS, AKT, CCC, GS, ND. Data curation: TC, EGK, TS, AKT, CCC, GS. Visualization: TC, EGK ND. Supervision: ND. Funding acquisition: ND, GS, AKT, CCC. All authors have read and agreed to the published version of the manuscript.

