## supplemental file for "Extracellular Vesicles Carry Distinct Proteo-Transcriptomic Signatures That are Different from Their Cancer Cell of Origin"

### Supporting Information

#### Patient recruitment and sample collection

The details of the IRB/oversight body that provided approval or exemption for the research described are given below: Institute Review Board approved protocols (GCO # 06-0996, 14-0318, and surgical consent)at the Department of Urology, Icahn School of Medicine at Mount Sinai, New York, 10029. All necessary patient/participant consent has been obtained and the appropriate institutional forms have been archived.

#### EV extraction from serum, urine, and cell culture medium using nanoDLD and ultracentrifugation

#### Cell culture EV isolation

Human HRPC cell lines, 22RV1, purchased from American Type Culture Collection (ATCC) and maintained in RPMI 1640 cell culture medium (GIBCO). 22RV1 cell lines are supplemented with 1% antibiotic and were monitored till 80-90% cell confluency was achieved. The supernatant was then extracted and centrifuged at 300 x g at 4◦C; the resulting cell pellet comprised of dead cells and cellular debris were then removed. The remaining supernatant is transferred into a new 50ml tube and further centrifugated at 2,000 × g, 4◦C for 30 minutes allowing for larger vesicles and remaining cell debris to be pelleted and removed. The supernatant is then transferred into another 50ml tube and diluted till the total volume is 3/4 of volume of the tube. Sequentially, the solution is centrifugated at 10,000 × g, 4◦C for 45 minutes followed by ultracentrifugation at 120,000 × g, 4◦C for 2 hours (using Beckman coulter, thick wall polypropylene tube, Cat # 355642). The pellet derived from the ultracentrifugation is washed and resuspended in PBS followed by another ultracentrifugation at 120,000 × g, 4◦C for 2 hours. Finally, the pellet is collected and resuspended in 1 ml of PBS and stored at -80C.

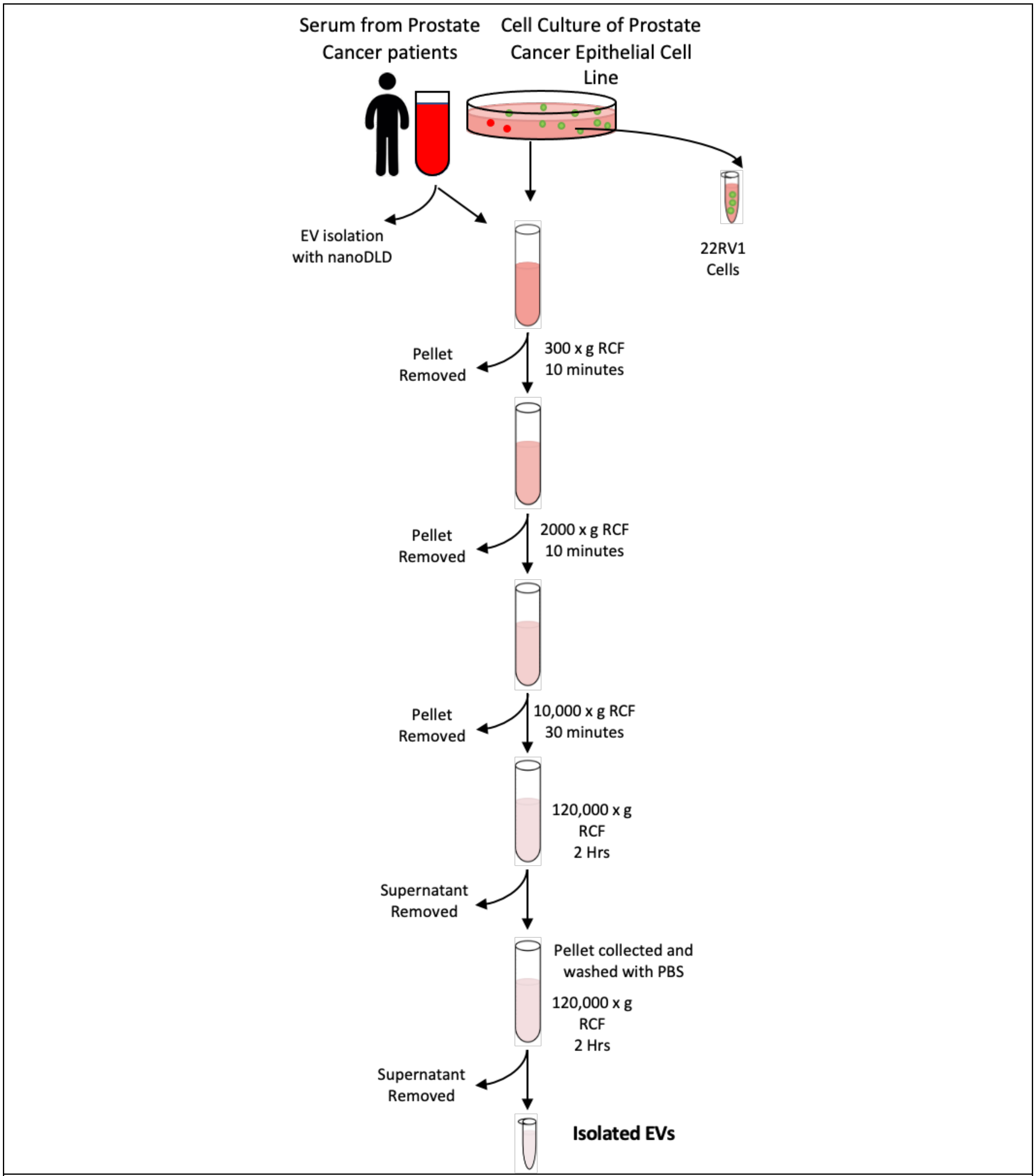


**EV isolation protocol**

### Serum derived EV isolation via ultracentrifugation

Blood from prostate cancer patients was collected via BD Vacutainer blood collection tubes and serum isolation was performed using serum separation tubes from Fisher Scientific (Cat.# 368016). 5 ml of the isolated serum aliquoted and centrifuged at 2,000 × g, 4◦C for 30 minutes. The supernatant is then transferred to a new 50 ml tube. To ensure the fluid volume is 3/4 of the total volume, PBS was added to the sample supernatant. The resulting solutions were centrifuged at 12,000 × g, 4◦C for 45 minutes. Sequentially, the supernatant is ultracentrifuged at 120,000 × g, 4◦C for 2 hours (using Beckman coulter, thick wall polypropylene tube, Cat # 355642). The pellet derived from the ultracentrifugation is washed and resuspended in PBS followed by another ultracentrifugation at 120,000 × g, 4◦C for 2 hours. Finally, the pellet is collected and resuspended in 1 ml of PBS and stored at -80C.

### Serum EV isolation via nanoDLD

Aside from the conventional methodology of EV isolation via ultracentrifugation, we have implemented an innovative method for serum EV isolation by utilizing the nanoDLD apparatus. To minimize non-specific adsorption, we have prepped the chips using a 0.02 µm-filtered solution of 5% (w/v) bovine serum albumin (Sigma Aldrich) in phosphate buffer saline. Samples were placed through the apparatus setting at G = 225 nanometer for 1 hour at Papp of approximately 5 bar. EVs in the range of 70-100 nanometers were primarily delegated into the bump fraction. Of note, smaller (<50 nm) particles remain in zigzag fraction of nanoDLD and are no collected for analyses. Rigorous characterization of UC and nanoDLD isolated EVs from various biofluids has been presented in our previous publications[1,2]

### Nanoparticle tracking analysis

Prior to performing nanoparticle tracking analysis, samples collected were further diluted using Millipore DI water to a targeted concentration of 106-107 particles/mL. ZetaView was then utilized to evaluate the particle size concentration and zeta potential via the built-in EMV Zeta protocol. Nanoparticle tracking analysis results verify the presence of EVs in the bump fraction of the serum samples from the nanoDLD apparatus. Likewise, the detected EV concentrations appeared to be ~2.6 to 3 times higher than that found in the input fractions.

### Immuno-fluorescence co-localization analyses (Nanoview)

Briefly, canonical tetraspanin exosome markers CD81, CD9, and CD63 against exosome surface are arrayed on silicon chips. EV suspensions are incubated with the chips overnight. After incubation, chips are washed with PBS on a shaker and air dried. Captured EVs are detected using Single Particle Interferometric Reflectance Imaging Sensor technology.

### specimen TEM analyses

### EV TEM analyses

EVs TEM analyses. Frozen EVEV pellet was brought to room temperature. Equal volumes of EVs and 3% Glutaraldehyde were mixed and kept at room temperature for 1 hr. Osmium tetraoxide was added to the EV solution and was kept at room temperature for 1 hr. The final EVs solution was transferred to formvar coated TEM grid. Observe under the electron microscope at 80 kV. Store the grids in the appropriate grid storage boxes for future use.

**Immuno-gold labeling of EVs**

Immuno-gold labeling of EVs. Frozen EVs pellet was brought to room temperature. Equal volumes of EVs and 3% Glutaraldehyde were mixed and kept at room temperature for 1 hr. 2ul of EV pellet was transferred to formvar coated TEM grid (at least 2 grids were prepared for each sample). TEM grids were covered and dried at room temperature for 30 minutes. 100ul drops of PBS were transferred to Parafilm. Dried TEM grids were carefully washed by transferring them on top of the PBS drops with the help of forceps (this step is repeated 3 times). The grid is transferred to a 100ul drop of BSA (this step is repeated 3 times). Then, the grid is transferred on top of a 5ul drop of primary antibody (CD81) in a blocking buffer for 30 minutes. Transfer the grid to a washing buffer/blocking buffer for 5 minutes (this step is repeated 5 times). A drop of 5ul gold conjugated secondary antibody is transferred to the Parafilm. The grid is transferred (and covered) on top of the gold antibody drop for 30 minutes. Once completely incubated, the grid is washed by keeping on top of 100ul PBS solution for 3 minutes (this step is repeated 10 times). Finally, contrast the EVs on the TEM grid with osmium tetraoxide for 10 minutes. The grid is ready for TEM imaging.

**RNA extraction, library preparation, and next-generation sequencing**

RNA Extraction, Library Preparation, and Next-Generation Sequencing. Total RNA was extracted from the serum bump fraction of nanoDLD, serum EVs pellet from UC using the Total EV RNA and Protein Isolation Kit (Invitrogen 4478545). 50 μL EVs from bump fraction, 100-200ul serum/urine EVs were resuspended in equal volume of ice-cold EV resuspension buffer. 2X denaturing solution was added to the final EVs solution on ice. Equal volume of acid-phenol:chloroform solution was added to each sample. The final solution was vortexed for 60 seconds and centrifuged at 10,000 x g. The top aqueous phase was carefully isolated without disturbing the lower organic phase. The top aqueous phase was transferred to the provided filter cartridge in collection tubes. Bound RNA was washed 3 times using the included wash solution. Finally, a preheated elution solution was used to elute the RNA in 100 μL. RNA was stored at -20 °C until it's RNA quality was assessed by bioanalyzer (Agilent 2100 Bioanalyzer, RNA 6000 Pico Kit, Agilent Technologies).

cDNA Libraries were prepared for small RNAs using the SMARTer smRNA-seq Kit for Illumina (Takara Bio 635030). A total of 18 cycles of PCR were carried out to obtain a good yield of cDNA from tissue, cells, and EVs. Final library quality was verified with Qbit and bioanalyzer. Negative (no RNA) and positive controls provided expected results. Next-generation RNA sequencing was performed using a HiSeq 4000 (Illumina), 100 base pair, single-end reads at the New York Genome Center.

**Mass Spectrometry**

Each frozen cell pellet was homogenized by adding pre-determined volume of lysis buffer (2%

SDS/1X protease inhibitor/0.1M Ambic). Enhanced BCA Protein Quantification assay was used to determine the total protein amount from each sample. Proteins from 30µg of lysates were separated from SDS using micro S-trap columns (http://www.protifi.com/s-trap/) and digested on column by trypsin. Resulting peptides were labeled with TMT6plex isobaric reagent per sample and then combined for high pH reverse phase peptide fractionation. Thermo Orbitrap Fusion Tribrid Mass Spectrometer was used for MS/MS analysis (MS3 data acquisition method). Three technical replications were run per sample. Proteins from 20ul of exosome lysates were separated from SDS using micro S-trap columns (http://www.protifi.com/s-trap/) and digested on column by trypsin. Resulting peptides were speedvac dried for LC-MS/MS analysis. Thermo Orbitrap Fusion Tribrid Mass Spectrometer was used for MS/MS analysis. Global normalization based on total number of ms/ms spectra (PSM) acquired was applied to the MS data. Spectral counts were used for semi-quantitative analysis to compare protein abundance among different samples.

Proteome Discoverer software (version 2.1) was used to search the acquired MS/MS data against a human protein database downloaded from the UniProt website and generate TMT ratios. Positive identification was set at 5% protein FDR and 1% peptide FDR. Also, at least 1 unique peptide has to be identified per protein. A total of quantifiable proteins from this study is 5630 proteins. TMT ratios (each tag/common reference) were calculated by PD 2.1 and normalized by total peptide amount. Qlucore Omics Explorer package was used to perform statistical analysis. Ingenuity Pathway Analysis (IPA) was used to perform data mining.
Proteome Discoverer software (version 1.4) was used to search the acquired MS/MS data against a human protein database downloaded from the UniProt website. Positive identification was set at 5% protein FDR and 1% peptide FDR. Also, at least 2 unique spectra has to be identified per protein. Scaffold Proteome Software was used for post-database search processing. 422 proteins passed the filtering criteria and their expression profiles among these four samples were analyzed to identify differentially expressed proteins. Qlucore Omics Explorer Statistical Software was used to perform appropriate statistical analysis.

**Computational Analyses**

Genome mapping:
For quantification of gene expression, raw reads were aligned to the latest Ensembl GRCh38.p13 (GCA_000001405.28) using bowtie aligner (version 2.5.4b). FeatureCounts was then used to map the aligned reads to the GENCODE v26 primary gene annotation, including transcripts corresponding to ncRNAS such as lncRNA, miRNA as well as protein-coding RNA. To maximise recovery and minimize the noise, multimapping reads were quantified up to m=10 and distributed using unique reads mapping distribution, as described in most recent best practices protocols.

Formal analysis:
Data cleaning, filtering, and analysis were performed in R and underexpressed genes or proteins with low or no counts across all samples of the similar phenotype were removed (at least one of the samples have CPM > 10). Normalization via trimmed mean of M-values in edgeR ensures library sizes of all samples are scaled properly to minimize the influences of external factors. The limma package, originally designed for microarray data, performs linear modeling on normally distributed data. Thus, to accommodate for the nonindependent mean-variance relationship of RNA-seq data, the voom function assigns a precision weight derived from the library size and

normalization factor of each sample itself to convert the raw counts to log2-CPM values. The log2-transformed counts minimize the changes in variance as the count size increases. Prior to examining differential expressions, we performed unsupervised clustering of samples to evaluate the similarities and dissimilarities between samples as well as across phenotypes of interest using the prcomp package in R. The result is reflected in the PCA plots.

Differentially expressed genes are discerned between 1) 22RV1 cell lines versus 22RV1 cell-line-derived EVs, and 2) Prostate cancer patient serum-derived EVs isolated using nanoDLD versus the EVs isolated using UC via the standard differential expression pipeline as illustrated in limma/edgeR packages. Results of the differentially expressed genes are represented in high-resolution heatmap as well as volcano plots made using pheatmap and ggplot2 packages. Likewise, differentially expressed proteins are discerned between 1) 22RV1 cell-line, 2) the EVs derived from the corresponding cell lines, and 3) patient serum EVs derived from UC versus those derived from nanoDLD.

Correlation analyses: Spearman Rho correlations were determined across cellular and EV genetic profiles as well as the proteomic profiles. Gene expressions were plotted in the x/y axis, where x/y axis are log2 (CPM), all RNA types were analyzed.

Biotype analysis: The gene biotype was recovered from the GTF annotation file for Ensembl GRCh38 (same as for aligment). Mapping resolution was kept as CDS with intron and exon annotation levels and combined to gene level when necessary. After differential expression quantification of gene biotype proportions, numbers and expression levels was taking into account. Thus, expressing gene biotype as (1) number of molecules per biotype (after lib. size adjustment) and (2) levels of expression using RPKM to adjust for gene/transcript length sizes.

Pathway Analysis: To effectively compare, not only enriched genes, but also against enriched proteins, pathway enrichment analyses are performed using the enrichR package in R. Specifically, we referenced databases including Kyoto Encyclopedia of Genes and Genomes (Versions: 2013, 2015, 2016, 2019, 2021), Gene Ontology Molecular Function (Versions: 2013, 2015, 2017, 2017b, 2018, 2021), Gene Ontology Cellular Component (Versions: 2013, 2015, 2017, 2017b, 2018, 2021), Gene Ontology Biological Process (Versions: 2013, 2015, 2017, 2017b, 2018, 2021), Reactome (Version: 2016), and WikiPathways (Version: 2019). Top 1500 enriched genetics and all of the proteomic signatures were used for the pathway analysis. Top 10 genomic and proteomic pathways from each database with FDR below 0.05 and at least three enriched genes present were selected. The overlaps across different cellular and EV datasets are outlined in the barplot made using the ggplot2 package in R. Similarly, we also performed the gene set enrichment analysis using our RNAseq result for hallmark cancer pathways. GSVA scores are generated per sample using the GSVA program in R. Wilcoxon test is then performed to identify significant GSVA scores across both cell and EV samples with a cutoff of 0.05 FDR. Of which, pathways with GSVA score differences greater than 0.1 across the cell and EV samples are then represented in a heatmap.

Finally, upon the acceptance of manuscript our data and results will be uploaded to GEO and will be openly available.

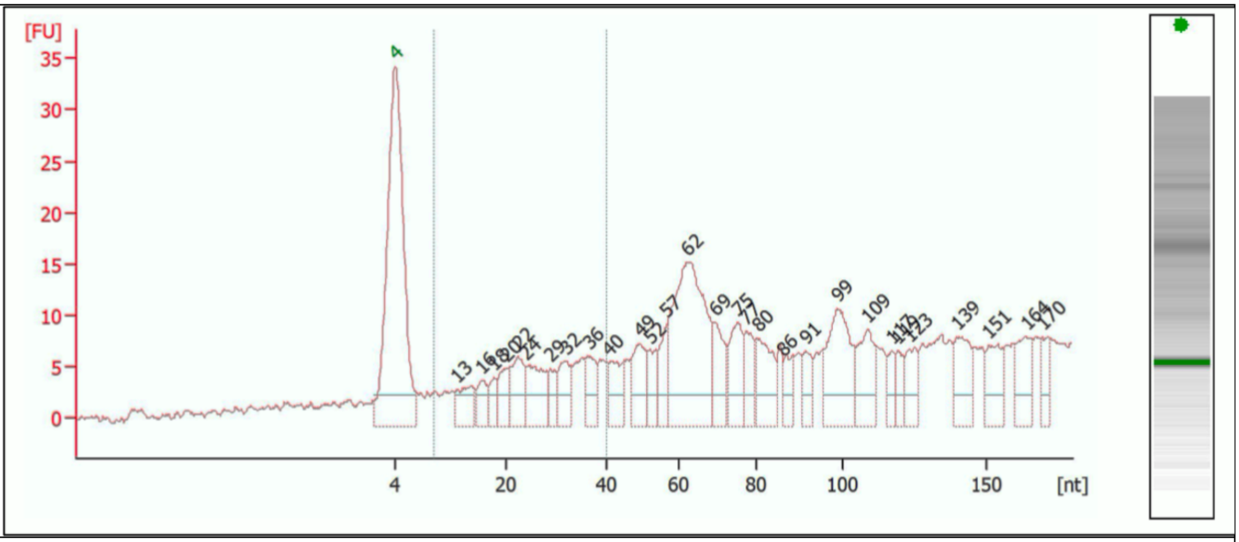


Supporting figure 1. Small RNA analyses of EV RNA show diverse RNA types ranging from 13-200 nt.

| 204 UC | 204 nanoDLD | 22rv1 EVs UC | interacting p | GO |
|---|---|---|---|---|
| MT-RNR2 | TCTE1 | VTRNA1-2 | IGF2BP2, SR: | mRNA bindir |
| MT-TV | AC127164.1 | VTRNA1-1 | present | |
| MIR126 | AC109466.1 | RNY1 | present | |
| MT-TP | PEX5L | MYOCOS | | |
| MIR25 | CCDC96 | RN7SL2 | | |
| MT-RNR1 | FLNA | SPEG | let-7 | srsf2 |
| MT-TG | CELF2 | AC124068.2 | | |
| RNY4 | SLPI | DPP9 | snhg16 enriched in evs | |
| MT-TD | ADGRG6 | VTRNA1-3 | igf2bp2 | |
| VTRNA1-1 | KCNIP2 | RNY4 | | |
| MT-TT | NRXN3 | RNY3 | ybx1 | TF |
| MT-TL2 | PLCB1 | MIR4454 | | |
| MT-TS2 | DBN1 | MAP1LC3B | | |
| MT-TL1 | WDR62 | AL049796.1 | | |
| MIRLET7G | ADRA1B | RNU5B-1 | | |
| MIRLET7G | UBLCP1 | LCN10 | | |
| MT-TR | AMZ1 | MEIS1-AS3 | | |
| MT-TH | MT-RNR2 | SMAD6 | | |
| AC124068.2 | ACSS3 | HES7 | | |
| RN7SL2 | GAB2 | BMPER | | |
| MIR23A | MCF2L | EHBP1L1 | | |
| MT-TM | ARFGAP2 | RNU5A-1 | | |
| RNY1 | SDCBPP2 | FBXO45 | | |
| HBB | CTC1 | POLR3A | | |
| MIR191 | GPR68 | RECQL4 | | |
| RNY3 | NUBP2 | Y_RNA | | |
| AL049796.1 | FMNL3 | P3H4 | | |
| MIR93 | TTPAL | RNA5SP426 | | |
| MIRLET7D | AC124068.2 | RPS6KA5 | | |
| HBA1 | ZNF25 | LENG8 | | |
| MIRLET7I | ZNF584 | LINC01287 | | |
| TCTE1 | KCNH4 | BPTF | | |
| MIR150 | AL032819.2 | CLIP2 | | |
| AC109466.1 | C4orf50 | YEATS2-AS1 | | |
| MT-TI | INSR | FDPS | | |
| HBA2 | USP15 | CIDEC | | |
| AC127164.1 | PSMD8 | IGFBP3 | | |
| PEX5L | NPFFR1 | GRINA | | |
| SLPI | FGFR1OP | DOC2B | | |
| CCDC96 | RNH1 | RNVU1-7 | | |
| PRKCB | SLC20A1 | AL355103.1 | | |
| MIR185 | DGKQ | VTRNA3-1P | | |
| MT-ND2 | MYO1C | HCN1 | | |
| CELF2 | AC095350.1 | FCGR2A | | |
| MT-TF | INF2 | TOR3A | | |
| MIR15B | RNA5-8SP5 | DLX2 | | |
| MT-ND4 | AC104457.1 | POU4F2 | | |
| FLNA | KLHL33 | FAM163A | | |
| MYOCOS | ZNF792 | LINC00324 | | |
| DBN1 | COL23A1 | ELN | | |

Table 1: Top 30 enriched RNA in EVs and complementary proteins for top 10 RNA.

1 Smith, J. T. *et al.* Integrated nanoscale deterministic lateral displacement arrays for separation of extracellular vesicles from clinically-relevant volumes of biological samples. *Lab Chip* **18**, 3913-3925, doi:10.1039/c8lc01017j (2018).
2 Murillo, O. D. *et al.* exRNA Atlas Analysis Reveals Distinct Extracellular RNA Cargo Types and Their Carriers Present across Human Biofluids. *Cell* **177**, 463-477.e415, doi:10.1016/j.cell.2019.02.018 (2019).
